# Metabolic rate and foraging behaviour: A mechanistic link across body size and temperature gradients

**DOI:** 10.1101/2024.03.02.583052

**Authors:** Milad Shokri, Francesco Cozzoli, Alberto Basset

## Abstract

The mechanistic link between metabolic rate and foraging behaviour is a crucial aspect of several energy-based ecological theories. Despite its importance to ecology however, it remains unclear whether and how energy requirements and behavioural patterns are mechanistically connected. Here we aimed to assess how modes of behaviour, in terms of cumulative space use, patch selection and time spent in an experimental resource patchy environment, are influenced by the foragers’ metabolic rate (SMR) and its main determinants i.e. body mass and temperature. We tested the individual behavioural patterns and metabolic rates of a model organism, the amphipod *Gammarus insensibilis*, across a range of body masses and temperatures. We demonstrated that body mass and temperature exert a major influence on foraging decisions and space use behaviour via their effects on metabolic rates. Individual cumulative space use was found to scale allometrically with body mass and exponentially with temperature, with patch giving-up time falling as body mass and temperature increased. Moreover, SMR had greater predictive power for behavioural patterns, explaining variation beyond that accounted for by body mass and temperature combined. Our results showed that cumulative space use scaled positively with Mass- and-Temperature-independent SMR (residual). Furthermore, the foraging decisions regarding patch choice and partitioning were strongly related to M-T independent SMR; individuals with higher M-T independent SMR initially preferred the most profitable patch and, as time progressed, abandoned the patch earlier compared to others. Our findings regarding the mechanistic relationship between behavioural patterns and metabolic rate across body mass and temperature shed light on higher-order energy-based ecological processes, with implications in the face of climate change.

## Introduction

Individuals differ considerably in their patterns of resource use in a heterogenous environment (Auer et al. 2020a), which has crucial evolutionary and ecological implications as it influences individual life history and space use, as well as population dynamics and community structure (Basset 1995, Auer et al. 2020b, Laskowski et al. 2022). Despite its importance to the higher-order ecological phenomena, the mechanisms underpinning individual behavioural variability remain the subject of active research debate.

At the fundamental level, foraging behaviour aims to fulfil organisms’ energy requirements (MacArthur and Pianka 1966). It is thus intuitively reasonable that metabolic rate, i.e. the baseline energy requirements of an organism (Brown et al. 2004), might be linked to foraging behaviour and resource acquisition strategies. Metabolic rate increases allometrically with organism body mass (Brown et al. 2004), and motile foragers need to adjust their space and resource use to fulfil their mass-dependent needs (Brose, 2010; Petchey et al., 2008). For example, larger foragers generally need larger home ranges and exploit a resource patch only if and for as long as it allows high ingestion rates (Basset, 1995; Charnov, 1976; Holling, 1959; McNab, 1963). In addition to individual body mass, as temperature increases, the higher kinetic energy of biochemical reactions speeds up the rate of metabolic processes (Gillooly et al. 2001), affecting the behavioural patterns of individual organisms (Abram et al. 2017). It follows that animals might adjust their foraging strategies in accordance with their energy requirements in response to body mass, ambient temperature or both (Dell et al., 2011; Petchey et al., 2008). Several studies have investigated the link between individual energy requirements and foraging behaviour, often focusing on one of the main determinants of metabolic rate, e.g. individual body mass (Brose, 2010; Cozzoli et al., 2019, 2022; Petchey et al., 2008) or, more recently, temperature (Abram et al. 2017, Cloyed et al. 2019, Jermacz et al. 2020). However, despite the key importance of body size and temperature *per se* in predicting behavioural patterns (Brose 2010, Cozzoli et al. 2022), they are not sufficient to crystalize the mechanism behind them (Brandl et al. 2022). The complexity of resource-consumer interactions often hinders the predictive power of mass and temperature-based models (Dell et al. 2014b). For instance, individuals may react to increased energy requirements by increasing foraging efforts and nutrient intake/ingestion (Réveillon et al. 2022) or, conversely, by limiting their movements (Warne et al. 2019). This is because animals are generally characterized by a set of biological rates that respond differently depending on the conditions (Barrios-O’Neill et al., 2019; Glazier, 2015). Alternative energy management hypotheses have been developed to explain the interdependence between metabolic rate and behavioural patterns. The performance model predicts that metabolic rate and activity will correlate positively because metabolic rate reflects activity (Biro and Stamps 2010). However, (Careau and Garland 2012) suggested that there is not always a positive correlation between metabolic rate and behavioural patterns. The allocation model assumes a negative relationship between metabolic rate and behavioural patterns, as a higher metabolic rate results in less energy available to spend on active behaviours such as locomotion and space exploration (Careau et al. 2008). The independent model sees individual behavioural activity and metabolic rate as independent (Careau and Garland 2012).

No consensus has emerged because metabolic syndromes cannot be completely understood without considering the effects of subsidiary environmental factors (Killen et al. 2013) and because behavioural studies have applied sharply differing methods, hampering comparative analysis (Montiglio et al. 2018). Additionally, investigating behavioural patterns in landscapes with homogenously distributed resources (or no resources at all) may mask the individual behavioural decisions related to energy requirements (Spiegel et al. 2017, Cloyed and Dell 2019). Here we aimed to investigate the interdependency across size and temperature gradients between metabolic rates and modes of behaviour (in terms of cumulative space use, patch choice and time spent) in a resource-patchy environment. Given the temperature-dependence of ectotherm metabolic rates, the incorporation of temperature into behavioural patterns via a metabolic framework can be expected to shed light on the responses to ongoing climate change of consumer-resource and predator-prey interactions.

To reduce the influence of confounding environmental factors, we performed controlled microcosm experiments using the aquatic amphipod *Gammarus insensibilis* as a model organism across a wide range of body masses and temperature regimes. Individual behavioural patterns were remotely quantified by using advanced automated image-based tracking (Dell et al. 2014a), and the standard metabolic rate of the specimens was accurately measured with an open-flow respirometry system (Glazier & Sparks, 1997). The factorial design allowed us to disentangle the link between standard metabolic rate (SMR) and the descriptors of time and space use behaviour across body mass (M) and temperature (T) gradients. To our knowledge, this is one of the first studies (see also (Réveillon et al. 2022) of multi-trait resource and space use behaviour accounting for factorial interactions among M, T and SMR under highly controlled microcosm conditions.

## Materials and methods

### Experimental design

The experiment made use of a full factorial design to assess the co-variation of individual foraging behaviour with SMR in an aquatic ectotherm model organism, i.e. *Gammarus insensibilis*, across a range of body masses and temperatures. The temperatures were selected to match the seasonal range experienced by *G. insensibilis* in its local environment. The water temperature of the collection site was monitored weekly from 2015 to 2019, and the assessment temperatures were *i.* the average winter temperature (13°C), *ii.* the average annual temperature (18°C) and *iii*. the average summer temperature (25°C). At each temperature level, we tested model organism specimens in a range of body sizes corresponding to the species’ size range in nature. The individual behaviour of *G. insensibilis* was assessed using automated video tracking while foraging alone in a heterogeneous environment under controlled microcosm conditions. The experimental microcosm (maze) was made up of six interconnected patches, two of which contained differing amounts of resources, i.e. a rich patch (1 g) and a poor patch (0.5 g), and four were empty, thus creating a heterogenous environment. After assessing foraging behaviour, individual SMR was measured using an open-flow respirometer system.

### Model organism

*Gammarus insensibilis* (Stock 1966) is a widespread Atlantic-Mediterranean amphipod species living in coastal and transitional waters, reaching maturity at 0.4 cm and growing to a maximum length of ∼2 cm (Costello et al. 2001). *Gammarus* species are selective foragers, feeding preferentially on fungi with high protein content colonizing decomposed leaf litter (Bärlocher and Kendrick 1973). While foraging they scroll the surface of the leaves seeking valuable resources, and when high-quality resources are not available, they consume resources with a lower value (Cozzoli et al. 2022). Daily consumption rates are from 46% to 103% of their body mass, depending on the type of exploited resource (Berezina 2007). Additionally, when resources are abundant, they have a gut throughput time of 45-59 minutes at 14°C (Welton et al. 1983). For the purposes of this study, the amphipod *G. insensibilis* is an ideal model organism due to the ease of manipulation in the laboratory and the recent development of experimental protocols (Shokri et al. 2021).

### Specimen collection and acclimation

Specimens of *G. insensibilis* were collected from the Cesine nature reserve, a transitional water body in Italy, at the southern end of the Adriatic Sea (40.3624 N, 18.3325 E). Authorization for specimen collection was issued by the competent authority (WWF, World Wildlife Fund for Nature, Italy). After collection, the specimens were transferred alive to the laboratory in thermo-insulated containers filled with water from the sampling sites and aerated during transport. The specimens were maintained in the laboratory aquaria at a salinity of 7 g l^-1^, similar to that of the sampling site, and acclimated to each of the assessment temperatures for two weeks in order to reduce the risk of temperature shock that might affect individual metabolic rates (Semsar-kazerouni and Verberk 2018). Acclimation to the low and high temperatures with respect to that of the sampling site at the moment of collection was achieved gradually at a rate of ±1.5 °C day^-1^ in aquaria under controlled climatic conditions (KW apparecchi scientifici, WR UR). Prior to the experiment, specimens were sorted by sex under a Nikon stereoscope (SMZ1270). Only adult males were selected for behavioural assessment, since oocyte production in females may induce non-size-related variability in energy requirements and behaviour, and to reduce potential variation due to different ontogenic stages (Glazier et al., 2011).

### Trophic resource preparation

Leaves of *Phragmites australis* (Cav.) Trin. ex Steud were collected at the specimen collection site, cut into approximately 10 cm lengths, oven dried at 60 °C for 72 hrs, weighed into separate portions (1 g for the Rich patch and 0.5 g for the Poor patch) and placed in 5 mm mesh plastic bags. The amount of resource in each patch (1 g, 0.5 g and 0 g) was considered to be sufficiently different for animals to distinguish between them (Cozzoli et al. 2022). The leaves were then leached and conditioned for two weeks in running environmental water at 18°C. The nutritional quality of the leaves is known to increase during conditioning because of microbial colonization and the assimilation of nutrients from the water (Boling et al. 1975).

### Foraging behaviour setup and measurements

The experimental system consisted of a microcosm (maze) made of transparent Plexiglas installed in an isolated and temperature-controlled room (KW apparecchi scientifici, WR UR). The microcosm was composed of six circular patches [13 cm in diameter, 3 cm high], connected by a network of channels [2.5 cm wide, 3 cm high] (Fig. 1b). The microcosm (maze) was placed on top of a near infrared backlight source in order to achieve high contrast, which facilitated specimen detection. Three infrared-sensitive cameras (Basler, aca1300-60gm) were mounted above the microcosm to film individual movement and patch use. The temperature treatments were evenly spaced throughout the experimental period to minimize possible varying acclimation effects.

**Figure 1.**
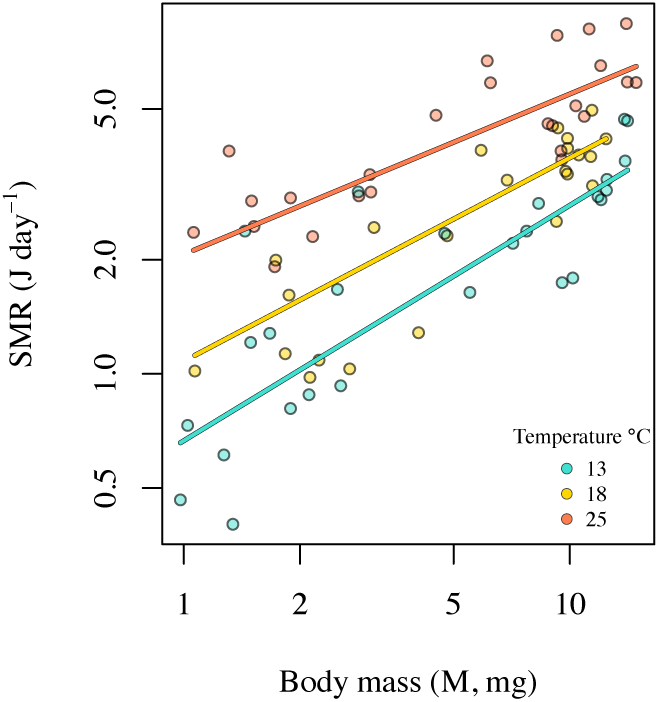
Standard metabolic rate (SMR, J day^−1^) in relation to body mass (M, mg) across temperature levels.

Prior to the behaviour assessment, each of the analysed specimens was kept unfed for 24 hrs in the climate-controlled room at the assessment temperature. This served to standardize specimens’ resource requirements at the start of experimental trials. For each experimental trial, 1 g dry weight of conditioned leaf fragments was placed in one patch and 0.5 g dry weight of conditioned leaf fragments was placed in another patch, thereby simulating a heterogenous resource distribution with two resource patches, “Rich” and “Poor”, while the other four patches were “Empty”. The distribution of the resource patches was randomized for each experimental trial to prevent any effect of microcosm geometry. The resource patches were placed in the microcosm 30 min before starting the experiment. Each experimental trial was performed on a single specimen foraging alone in the microcosm. The experimental trials were always conducted at the same time of day (09:00 to 15:00) to prevent any effect of the model organism’s circadian rhythms. Recordings were initiated 10 min after the specimen was released into the microcosm and lasted for 6 hrs. The video files were then processed by Ethovision XT 14 in batch acquisition mode, with the specimens identified by the software as moving elements with respect to the static background. A patch was considered to have been “visited” once the specimen had travelled the full length of a channel, entered a neighbouring patch and stayed there for at least 30 seconds.

### SMR and body mass measurements

After the behavioural measurement, the specimens’ Standard Metabolic Rate (SMR, J day^-1^) was measured. Although the sequential arrangement of metabolic measurements following behavioural observations might hold the potential to introduce a degree of uncertainty, specimens were kept unfed individually for 24 hrs before the SMR measurements. This step was taken to standardize the conditions, minimize residual effects from the behaviour experiment, and also ensure the specimens were in post absorptive state, as 24 hrs is sufficient to complete digestion in *Gammarus* sp. (Welton et al. 1983). Following (Glazier & Sparks, 1997), individual Standard Metabolic Rate (SMR, J day^-1^) was determined by measuring the oxygen consumption. Animals were placed individually in Strathkelvin open-flow system respirometers where the oxygen concentration was continuously measured by Clark-type microelectrodes connected to an oximeter and recorded using the Strathkelvin software (SI, 929). A 0.3 mm nylon mesh with a nominal outer diameter of 12.07 mm was placed in each respirometer chamber in order to minimize the individual’s spontaneous movement. After metabolic measurement, the animals were dried individually in an oven at 60 °C for 72 hrs and then weighed on a micro balance (Sartorius MC5) to the nearest ± 0.001 mg.

### Data analysis

The scaling of individual standard metabolic rate (SMR, j day^-1^) with individual body mass (M, mg) and temperature (T) was assessed via multiple linear regression. The response variable individual SMR and the explanatory variable M were log-transformed in order to fit the size-scaling relationship as a power law (Brown et al. 2004), and the temperature was inverse transformed to linearize its effect (Brown et al., 2004; Gillooly et al., 2001):

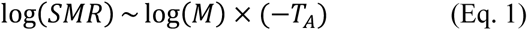

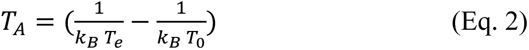

*T*_A_ is a standardised inverse temperature, *k_B_* is the Boltzman constant (8.618×10^-5^ *ev*/*k*), *T_e_* is the assessment temperature, and *T_0_* sets the intercept of the relationship at 286.15 K, corresponding to the lowest temperature level (i.e.,13 °C in this study). Standardising the inverse temperature at *T_0_*, simplified the interpretation of the main effect coefficients in the presence of interactions. The multiple linear regression was fitted with full interaction between explanatory variables.

We analysed the specimens’ behavioural patterns in the experimental microcosm with reference to three descriptors of space and time use behaviour. (1) cumulative space used, approximated as the total number of patches visited or revisited during the experiment. For the temporal trends in patch use throughout the experimental time (i.e. 360 min), we quantified (2) the proportion of specimen’s time spent in patches with resources (rich and poor) within each 5-min time interval, and (3) the proportion of time spent in the Rich patch relative to the time spent in patches with resources within each 5-min time interval. Descriptor (2) describes the temporal trends of the specimens’ time spent in resource patches compared to empty patches, and descriptor (3) reflects how the specimens allocated their time between resource-rich and resource-poor patches. A higher value indicates more time spent in the Rich patch, whereas a lower value shows more time spent in the Poor patch by the specimens.

Individual variation in the behavioural descriptors was analysed across individual body mass (M), temperature (T), and M and T independent SMR (residual SMR) gradients. We estimated the SMR residuals, the components of a metabolic phenotype that is independent of mass and temperature, using Eq. 1. This is hereafter referred to as M-T independent SMRs. We used this approach to examine whether SMR affects behavioural patterns beyond the combined effects of size and temperature.

The variation in descriptor (1) (i.e. the cumulative number of patches visited during the experiment (N)) was investigated by linear regression along the M, T, and the M-and T-independent SMR gradient. Both the response variable N and the explanatory variable M were log-transformed. As with the previous analysis (Eq. 1), the explanatory variable temperature was inverse transformed. The multiple linear regression was fitted with full interaction between explanatory variables.

For both descriptors of the temporal trend in patch use (2 and 3) we conducted binomial Generalized Linear Mixed Models (GLMMs) with a logit link. For descriptor (3) the model accounted for trial binomial data, wherein each observation represents the number of successes out of a given number of trials (Douma & Weedon, 2019). This was chosen to handle the varying denominators associated with the proportions of time spent in the Rich patch relative to patches with resources. For both of temporal trends in patch use descriptors, the models were defined as a function of the continuous explanatory variables M, T, and M-T independent SMR along with the experimental time. The explanatory variable forager body mass was log-transformed in order to model size dependency as a power law and the temperature was inverse transformed. In our models assessing temporal trends in patch use, to account for the non-independence of observations repeated over time on the same individual, we initially fit the models with both random intercepts and slopes at the individual level. However, the variance attributed to the random slope for time was negligible (τ11 in all models was <0.0001). Given the minimal contribution of the random slope and to avoid potential model overcomplication, we allowed random variation in the intercepts at the individual level.

The relative importance of mass, temperature, and M- and T-independent SMR in explaining the variance of the response variable was assessed by the LMG metric (R^2^ partitioned by averaging over orders (Lindeman et al. 1980)). The uncertainty of model estimates was reported as the 95% Confidence Interval [lower-upper]. All analyses were performed within the ‘R’ free software environment (R Core Team 2019) using the lme4 (Bates et al. 2015), relaimpo (Groemping 2006), partR2 (Stoffel et al. 2021) and sjPlot (Lüdecke 2018) packages.

## Results

### Specimen characterization

The 75 male specimens of *G. insensibilis* used in this experiment ranged from 4.73 to 19.86 mm in body length (on average 11.55 mm [± 4.04 SD]) and from 0.98 to 14.85 mg dry weight in body mass (on average 6.59 mg [± 4.46 SD]). The body mass distribution of the analysed specimens was similar across the three temperature treatments (ANOVA; F_2,72_ = 0.07, p = 0.93).

### Size scaling SMRs across temperatures

Overall, specimens’ individual Standard Metabolic Rates (SMR) ranged from 0.40 to 8.39 J day^-1^ (on average 3.15 J day^-1^ [± 1.81 SD]) and increased with temperature, with an average of 2.09 J day^-1^ [± 1.19 SD] at 13 °C, 2.79 J day^-1^ [± 1.28 SD] at 18 °C and 4.57 J day^-1^ [± 1.90 SD] at 25 °C. 76.1 % of the variation in individual SMRs was explained by the positive dependency on M (48% of explained variance, scaling exponent 0.62 [0.50 – 0.74 95% CI]) and T (28.1% of explained variance, scaling exponent 0.69 [0.48 – 0.91 95% CI]) (Table 1, Fig. 1). An additional 2.9% of the observed variance in SMR was explained by the marginal negative interaction between M and T (Table 1, Fig. 1), implying that the rate of increase of SMR with temperature decreased slightly as body mass increased.

**Table 1.** Output of the linear regression model of the variation in standard metabolic rate (SMR, J day^-1^) across body mass (M) and temperature (T) gradients.

| log (SMR) |  |  |  |  |  |
| --- | --- | --- | --- | --- | --- |
| *Predictors* | *Estimates* | *CI* | *t value* | *p* | *df* |
| (Intercept) | -0.40 | -0.61 – -0.19 | -3.81 | <0.001 | 71 |
| log (M) | 0.62 | 0.50 – 0.74 | 10.33 | <0.001 | 71 |
| T | 0.69 | 0.48 – 0.91 | 6.45 | <0.001 | 71 |
| log (M) × T | -0.12 | -0.24 – -0.00 | -2.03 | 0.046 | 71 |
| Observations | 75 |  |  |  |  |
| R2 / R2 adjusted | 0.791 / 0.782 |  |  |  |  |

### Space use

49.7% of the observed variation in foragers’ cumulative space use was explained by its positive allometric scaling with M (21.9 % of explained variance, scaling exponent 0.47 [0.27 – 0.67 95% CI], Table 2, Fig. 2a), the positive exponential relationship with T (27.8% of explained variance, exponent 0.72 [0.36 – 1.08 95% CI], Table 2, Fig. 2a). The interaction of M and T was not significant (Table 2). Additionally, cumulative space use scaled positively with the foragers’ M-T independent SMR, with a scaling exponent of 0.84 [0.44 – 1.24 95% CI] (Table 2, Fig. 2b). This explained an additional 10.1% of the variation in cumulative space use, beyond what was accounted for by body mass and temperature combined (Table 2, Fig. 2b). The significant relationship between cumulative space use and M-T independent SMR, along with its explained variation, indicates that SMR, when considered as a single descriptor in its entirety, explains a greater amount of variance than that explained by body mass and temperature combined.

**Figure 2.**
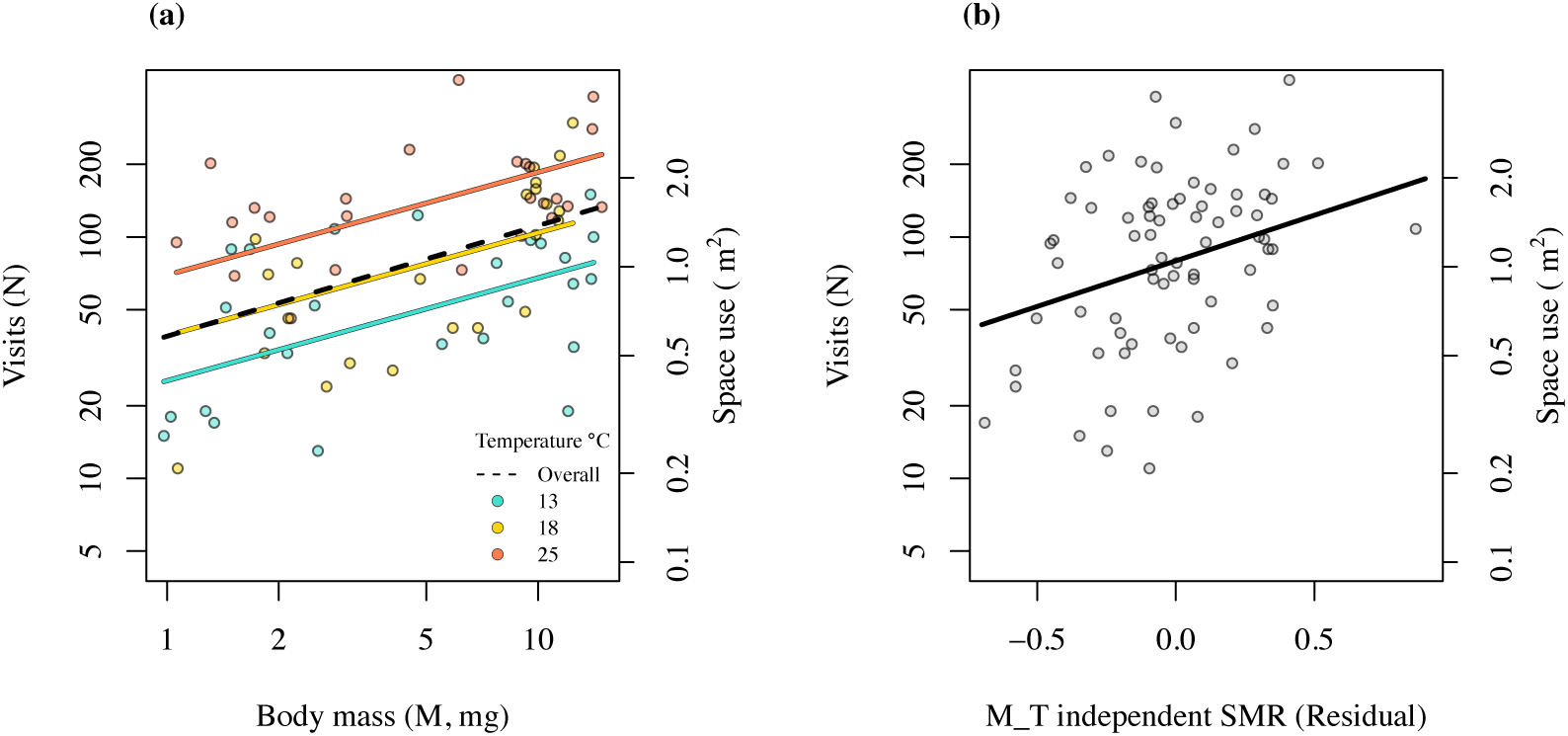
(a) Total number of visits to all patches in relation to body mass (M, mg) across temperature levels. (b) Total number of visits to all patches in relation to M and T-independent standard metabolic rate (residual SMR). Positive values of M-T independent SMR indicate individuals with higher-than-average metabolic rates for their size and the given temperature, while negative values indicate the converse. The secondary y-axis shows cumulative space use (m^2^), calculated as the overall surface area of patches that individuals visited.

**Table 2.** (a) Output of the linear regression between the total number of visits to all patches, i.e. cumulative space use, with respect to individual body mass (M), temperature (T) and M-T independent SMR (residual).

| log (total visits) |  |  |  |  |  |
| --- | --- | --- | --- | --- | --- |
| *Predictors* | *Estimates* | *CI* | *t value* | *p* | *df* |
| (Intercept) | 3.16 | 2.81 – 3.52 | 17.75 | <0.001 | 70 |
| log (M) | 0.47 | 0.27 – 0.67 | 4.64 | <0.001 | 70 |
| T | 0.72 | 0.36 – 1.08 | 3.96 | <0.001 | 70 |
| SMR (Residual) | 0.84 | 0.44 – 1.24 | 4.21 | <0.001 | 70 |
| log (M) × T | -0.06 | -0.27 – 0.14 | -0.63 | 0.528 | 70 |
| Observations | 75 |  |  |  |  |
| R<sup>2</sup> / R<sup>2</sup> adjusted | 0.601 / 0.578 |  |  |  |  |

### Temporal trends in patch use

At the beginning of the experiment, the specimens displayed a marked preference for the Rich patch, spending most of their time there, while the Poor and Empty patches were largely ignored (Table 3, 4, Fig. 3). The initial preference for Rich patch was pronounced at increased temperature (Table 4, Fig 3c), and was also observed among specimens with a higher M-and T-independent SMR (Table 4, Fig. 3d).

**Figure 3.**
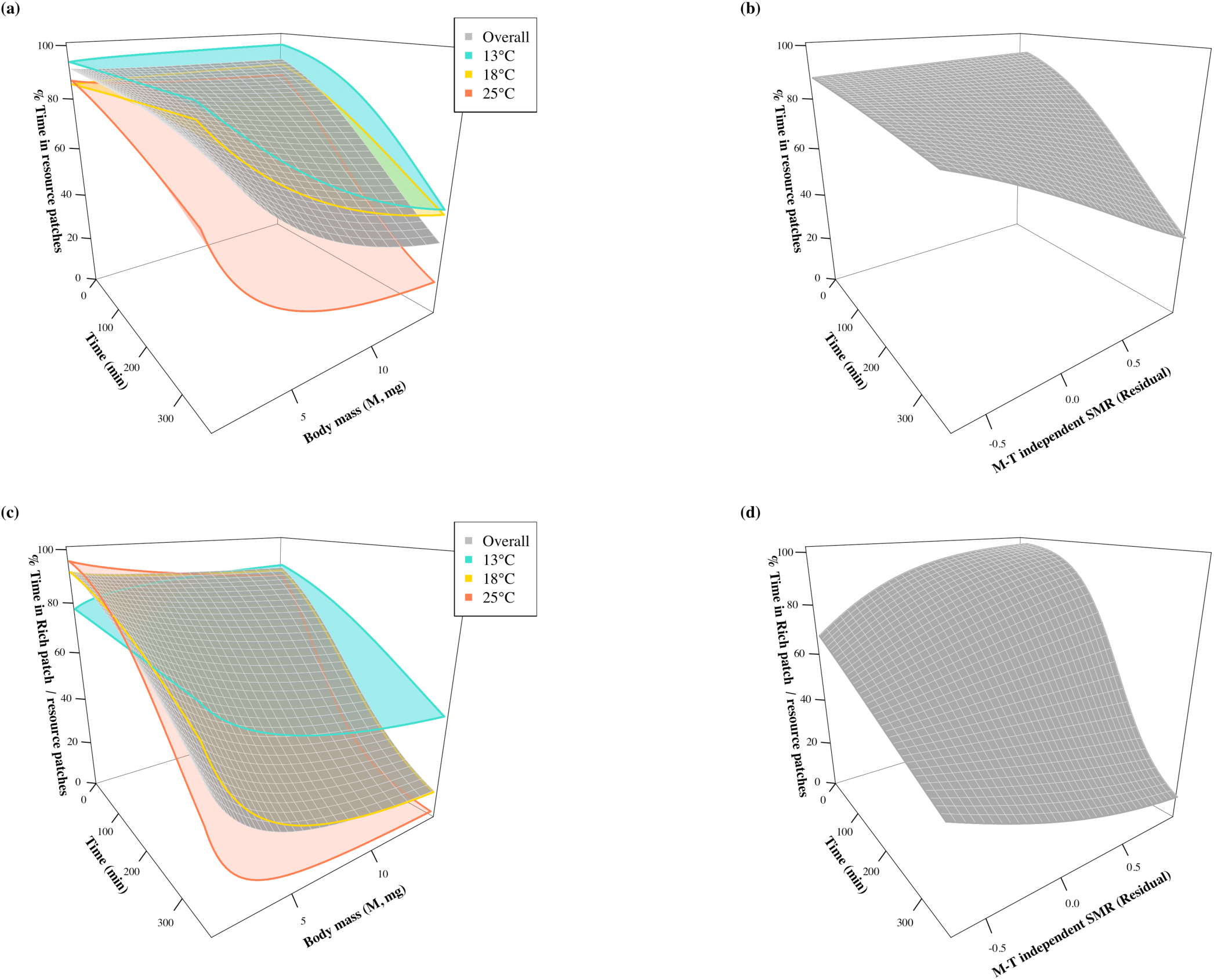
(a-b) Model surfaces of time spent (%) in patches with resources within 5 min time intervals (a) with respect to experimental time, body mass and temperature, and (b) with respect to M- and T-independent SMR. Positive values of M-T independent SMR indicate individuals with higher-than-average metabolic rates for their size and the given temperature and negative values indicate the converse. (c-d) Model surfaces of time spent (%) in Rich patch relative to resource patches (c) with respect to experimental time, body mass and temperature, and (d) with respect to M- and T-independent SMR.

**Table 3.** Output of binomial generalized linear mixed-effects model, where the response variable is the proportion of specimen’s time spent in patches with resources within the 5-minute time interval. The explanatory variables include body mass (M), temperature (T), M-T independent SMR (residual) and experimental time as the continuous explanatory variables.

| *Predictors* | %Time in resource patches |  |  |  |
| --- | --- | --- | --- | --- |
|  | *Odds Ratios* | *CI* | *z value* | *p* |
| (Intercept) | 14.40 | 6.29 – 32.95 | 6.31 | <0.001 |
| Time | 1.00 | 1.00 – 1.00 | 0.13 | 0.897 |
| log (M) | 1.08 | 0.69 – 1.69 | 0.34 | 0.735 |
| T | 0.53 | 0.24 – 1.15 | -1.61 | 0.107 |
| SMR (Residual) | 3.60 | 0.77 – 16.89 | 1.62 | 0.104 |
| Time × log (M) | 1.00 | 1.00 – 1.00 | -6.65 | <0.001 |
| Time × T | 1.00 | 1.00 – 1.00 | -2.46 | 0.014 |
| Time × SMR (Residual) | 0.99 | 0.99 – 1.00 | -5.56 | <0.001 |
| log (M) × T | 0.94 | 0.62 – 1.42 | -0.29 | 0.772 |
| log (M) × SMR (Residual) | 1.05 | 0.35 – 3.09 | 0.08 | 0.936 |
| T × SMR (Residual) | 0.58 | 0.15 – 2.26 | -0.79 | 0.432 |
| Random Effects |  |  |  |  |
| $\sigma^2$ | 3.29 | | | |
| $\tau_{00}$ individual | 1.07 | | | |
| ICC | 0.25 |  |  |  |
| N individual | 75 |  |  |  |
| Observations | 5400 |  |  |  |
| Marginal R<sup>2</sup> / Conditional R<sup>2</sup> | 0.241 / 0.422 |  |  |  |

**Table 4.** Output of binomial generalized linear mixed-effects model, where the response variable is the proportion of time spent in Rich patch relative to resource patches over 5-minute time intervals. The explanatory variables include body mass (M), temperature (T), M-T independent SMR (residual) and experimental time as the continuous explanatory variables.

| %Time in Rich / Resource patches |  |  |  |  |
| --- | --- | --- | --- | --- |
| *Predictors* | *Odds Ratios* | *CI* | *z value* | *p* |
| (Intercept) | 1.87 | 0.42 – 8.33 | 0.82 | 0.410 |
| Time | 1.00 | 1.00 – 1.00 | 3.03 | 0.002 |
| log (M) | 2.28 | 0.98 – 5.29 | 1.92 | 0.053 |
| T | 20.93 | 4.63 – 94.57 | 3.95 | <0.001 |
| SMR (Residual) | 28.49 | 1.49 – 543.73 | 2.23 | 0.026 |
| Time × log (M) | 1.00 | 1.00 – 1.00 | -10.46 | <0.001 |
| Time × T | 0.99 | 0.99 – 0.99 | -34.99 | <0.001 |
| Time × SMR (Residual) | 0.98 | 0.98 – 0.99 | -25.82 | <0.001 |
| log (M) × T | 0.31 | 0.14 – 0.72 | -2.73 | 0.006 |
| log (M) × SMR (Residual) | 3.55 | 0.38 – 32.91 | 1.12 | 0.264 |
| T × SMR (Residual) | 0.16 | 0.01 – 2.69 | -1.27 | 0.205 |
| Random Effects |  |  |  |  |
| $\sigma^2$ | 3.29 | | | |
| $\tau_{00}$ individual | 1.29 | | | |
| ICC | 0.28 |  |  |  |
| N individual | 75 |  |  |  |
| Observations | 5400 |  |  |  |
| Marginal R<sup>2</sup> / Conditional R<sup>2</sup> | 0.303 / 0.501 |  |  |  |

As the experimental time progressed, the time spent in the Rich patch decreased significantly, while it increased in the Poor patch (Table 3, Fig. 3c, d). The shift from time spent in the Rich patch to time spent in the Poor patch occurred earlier as body mass and temperature increased (Table 4, Fig. 3c). Moreover, this shift from the Rich towards the Poor patch occurred earlier in individuals with higher M-and T-independent SMR (Table 4, Fig. 3d). This implies that individuals with higher-than-average metabolic rates for their size and the given temperature, those having higher M-and T-independent SMR, were more prone to change their residency from Rich to Poor patch as experimental time continued.

Towards the end of the experimental time, the percentage of time spent by specimens with larger body mass and by all specimens at the higher temperature decreased in patches with resources (Table 3, Fig. 3a). Similarly, specimens with higher M-T independent SMR exhibited a decrease in patches with resources towards the end of the experimental time, corresponding to an increase in time spent in Empty patches (Table 3, Fig. 3b).

The fixed effects including M, T, and M-T independent SMR, explained a significant and comparable amount of variation in patch use behaviour (Table 3, 4). They collectively explained 24.1% of the variance in the proportion of time spent in patches with resources and 30.3% in the proportion of time spent in Rich patches relative to resource patches (Table 3, 4). Furthermore, the estimated random variation across specimens was 18.1%, for the proportion of time spent in patches with resources and 19.8% for the proportion of time spent in Rich patches relative to resource patches, highlighting the substantial individuality in patch use behaviour (Table 3, 4).

## Discussion

Overall, we observed that foragers modulate their resource and space use behaviour in response to variations in body mass, temperature and M-and T-independent SMR, highlighting the role of Standard Metabolic Rate (SMR) in its entirety as the key predictor of foraging patterns. This is likely because SMR (i) is mechanistically related to the individual’s energy balance and resource needs, rather than being a proxy of it, (ii) encompasses mass and temperature variations, (iii) is able to capture variations in energy needs beyond size and temperature, linked for example to life style and phenotype (Killen et al. 2010), and (iv) is intimately intertwined with informational control such as hormones (see Glazier, 2015).

### Size scaling SMRs across temperatures

In accordance with MTE expectations (*sensu* Brown et al., 2004), we observed that individual SMR increased allometrically with body mass and exponentially with temperature. However, the mass scaling exponents of SMR marginally decreased as temperature rose, implying that temperature-induced increases in metabolic rate are less pronounced in large-sized individuals than smaller ones. The latter observation accords with the Metabolic-Level Boundaries hypothesis (*sensu* Glazier, 2005; Glazier, 2020; Glazier, 2014) and with empirical evidence e.g. (Hoefnagel and Verberk 2015, Shokri et al. 2022) that the effect of temperature on metabolic rate is body mass-dependent.

### Space use

Individual cumulative space use was found to scale allometrically with body mass, implying that larger individuals used more space than smaller ones. This is consistent with the classical framework of (McNab 1963) and with empirical studies e.g. (Minns 1995, Cozzoli et al. 2022, Udyawer et al. 2022). On the other hand, we observed that individuals increased their space and resource use as a function of increasing temperature (within thermal tolerance), likely via kinetic effects (*sensu* Abram et al., 2017). Furthermore, we observed a marginal negative interaction between body mass and temperature in relation to SMR, but this was not reflected in the cumulative space use behaviour of the specimens. This discrepancy may indicate an adaptive behavioural response to warming. In response to temperature rise, organisms, especially the larger ones, face depletion of somatic energy resources from increased metabolic maintenance costs (see Glazier, 2015). As a result, they likely explored a greater cumulative space to access new resources and intensified their foraging effort to fulfill these demands.

We observed that individuals with a high SMR, after accounting for body mass and temperature, cumulatively explored a larger proportion of the space. This implies that individuals with higher M-and T-independent metabolic rates are able to collect, process and invest more energy and explore a greater space and resource in order to meet their requirements (since more energy would be needed to maintain this level of metabolism (see also Metcalfe et al., 1995)). The observed positive correlation between SMR and cumulative space use supports and extends the prediction from the performance model (*sensu* Biro & Stamps, 2010), demonstrating that higher metabolic rates require a larger area to explore for resource gathering in a patchy distributed environment.

### Temporal trends in patch use

Specimens at increased temperatures, as well as those with higher M-and T-independent SMR (beyond those dictated by body mass and temperature), exhibited a marked preference, spending more time in the Rich patch during the early hours of the experiment. This is likely because a more profitable patch offers a higher energy gain per unit of time, enabling foragers to achieve their optimal ingestion rate in a heterogeneous environment (MacArthur & Pianka, 1966; Stephens & Krebs, 1986). As the experimental time progressed, we observed that the time spent in the Rich patch decreased. This trend was sharper for larger foragers, at increase of temperature, and those with higher M-and T-independent SMRs. The specimens that left the resource-rich patch were observed to move and spend time in the Poor patch. Our observations on patch use align with the theoretical frameworks (MacArthur and Pianka 1966, Charnov 1976) and extend these concepts by empirically demonstrating that foraging decisions are intimately linked to an individual’s metabolic rate. This connection provides a mechanistic understanding of how metabolic processes influence the resource acquisition strategies, adding a new dimension to our understanding of foraging behaviour. The findings on these temporal trends of patch selection and use may potentially be explained by the combined effect of three mechanisms: (i) A higher SMR requires higher ingestion rates (Rosenfeld et al. 2015); foragers with higher SMRs thus deplete the resource patch more rapidly, resulting in shorter giving-up times than foragers with lower SMRs. (ii) Foragers with higher SMRs leave the patch when the amount of resource reaches a level (the marginal value (Charnov 1976)) that can no longer fulfil their energy requirement rapidly enough, even though it would still be economically viable for foragers with lower SMRs. This suggests that foragers with high metabolic rates are less able to exploit patches until reaching a low level of resource (or patches that are resource-poor to begin with), leading to a higher giving-up density than individuals with lower SMRs (see Cozzoli et al., 2018; Kotler et al., 1993; Kotler & Brown, 1990). (iii) Individuals’ resource specialization increases with energy requirement (e.g., intrinsically or with warming), because resources with higher energy content provide more energy per unit of processing time (Schoener 1974, Petchey et al. 2010). It follows that foragers with high SMRs exploiting a resource patch should also perceive a faster decrease in the available resource. Based solely on body mass, larger individuals are thought to be more selective of resources, as they have a higher total energy requirement per unit of time than smaller ones (Cozzoli et al. 2022). However, as temperature increased, speeding up metabolic rates, we found that individuals with a smaller body mass began to be more selective and perceive resource shortages in a similar way to larger ones at lower temperatures. This finding highlights the synergistic effects of body mass and temperature on resource specialization and perception via metabolic pathways. Moreover, while this observation supports the size dependency of foragers’ perception of available resources (*sensu* Basset et al., 2012; Basset & De Angelis, 2007), it extends this framework from body size to metabolic dependence.

In light of our findings and the urgent concerns over the effects of climate change, further studies of species and populations across latitudes under various climate change scenarios are needed. This would further our knowledge of these mechanisms and the adaptive behavioural responses to global warming.

### Ecological implications in the face of climate change

The mechanistic link between metabolic rates and resource and space use behaviour raises the prospect of understanding higher-order ecological processes, e.g. consumer-resource interaction, in the context of climate change. Under global warming scenarios, the equilibrium resource density is expected to decrease due to the temperature-driven rise in consumer metabolic and feeding rates being greater than any corresponding rise in resource growth and turnover rates (Bruno et al. 2015). This is expected to lead to stronger intraspecific competition among foragers owing to declining resource density, as well as increased top-down control in the food web (Lindmark et al. 2018). As a result of lower resource availability and increasing energy requirements with warming, animals are expected to range farther afield, leading to larger home ranges and niche overlap (Börger et al. 2008). This can expose animals to greater predation risk (Biro et al. 2009, Metcalfe et al. 2016, Balaban-Feld et al. 2022) and lead to stunted growth over time (Huey and Kingsolver 2019, Lackey and Whiteman 2022), as the organism may not be able to replace the energy it expends, because ingestion increases at a slower rate than metabolic rate as temperature rises (Basset et al. 2012, Réveillon et al. 2022).

In summary, this study offers insights that contribute to bridging metabolic and foraging theories. It showed that warming may have a profound influence on space/resource use and foraging decisions of individuals of different sizes, which have far-reaching consequences for higher-order ecological processes. Our results further highlighted the role of metabolic rate as the key predictor of behavioural patterns which encompasses variations that extend beyond those attributed to body mass, temperature, or their combination.

